# *Yersinia pestis* lipopolysaccharide remodeling confers resistance to a *Xenopsylla cheopis* cecropin

**DOI:** 10.1101/2021.03.12.435208

**Authors:** Basil Mathew, Kari L. Aoyagi, Mark A. Fisher

## Abstract

Fleas are major vectors of *Yersinia pestis,* the causative agent of plague. It has been proposed that *Y. pestis* has developed the ability to overcome the innate immune responses of fleas. Despite the fact that they transmit a number of bacterial infections, very little is known about the immune responses in fleas. In this study, we describe the antimicrobial activities of cecropin from *Xenopsylla cheopis* (cheopin), a major vector for *Y. pestis* in the wild. This is the first cecropin-class antimicrobial peptide described from Siphonaptera insects. Cheopin showed potent activity against Gram-negative bacteria, but little activity against wild-type *Y. pestis* KIM6+. Deletion of the aminoarabinose operon, which is responsible for the 4-amino-4-deoxy-L-arabinose (Ara4N) modification of LPS, rendered *Y. pestis* highly susceptible to cheopin. Confocal microscopy and whole cell binding assays indicated that Ara4N modification reduces the affinity of cheopin for *Y. pestis.* Further, cheopin only permeabilized bacterial membranes in the absence of Ara4N-modified LPS, which was correlated with bacterial killing. This study provides insights into innate immunity of the flea and evidence for the crucial role of Ara4N modification of *Y. pestis* LPS in conferring resistance against flea antimicrobial peptides.

## Introduction

The high virulence [1,2], recent outbreaks [3–6], emergence of multidrug resistant strains [7] and the presence of natural foci across all major continents, including Asia, Africa and Americas [8] draws a special attention to *Yersinia pestis,* the causative agent of plague. The natural reservoirs of *Y. pestis* in the wild are maintained by successful transmission of infections among rodents and other mammals through infected fleas, a group of insects belonging to the Order Siphonaptera. Fleas are the major vector of other significant zoonotic bacterial diseases, such as murine typhus *(Rickettsia typhi)* and cat-scratch disease *(Bartonella henselae)* [9]. Nonetheless, little is known about the innate immune responses of Siphonapteran insects [10].

Insects are known to produce a repertoire of structurally distinct groups of antimicrobial peptides (AMP) and AMP-mediated killing is a primary immune mechanism in insects [11–13]. *Xenopsylla cheopis,* the oriental rat flea, is a major vector for *Y. pestis* in nature [14–16]. Analysis of *X. cheopis* transcriptomes indicated that different classes of AMPs are expressed in fleas and are responsive to *Y. pestis* infections [17–19]. However, little is known about the antimicrobial activities or mechanisms of action of these peptides. Interestingly, *Y. pestis* has developed an arsenal of tools to overcome host immune responses [20]. The cell envelope of *Y. pestis* undergoes rapid remodeling and is proposed to be one of the key mechanisms by which *Y. pestis* successfully evade the host-immune responses [21–24]. At temperatures similar to that of the flea in the environment (21-28°C), phosphate groups of lipid A molecules are extensively modified with positively-charged 4-amino-4-deoxy-L-arabinose (Ara4N) [25–28]. This charge reversal is proposed to inhibit electrostatic interactions between cationic antimicrobial peptides and bacterial outer membranes (OM), leading to resistance against various classes of AMPs [29]. Incorporation of Ara4N into lipid A is primarily controlled by the PhoPQ two component system in *Yersinia* and other *Enterobacterales* [21, 28, 30]. Interestingly, in certain strains of *Y. pestis,* the presence of Ara4N on the LPS had negligible influence on the resistance against polymyxin B (PB), a prototypical cyclic AMP. Rather, a PhoPQ-mediated alteration of the oligosaccharide compositions of the LPS core confers resistance against PB in these strains [31]. Although there is no direct evidence of its effects on the antimicrobial activities of flea AMPs, *Y. pestis* with mutations in genes associated with Ara4N and PhoPQ pathways showed reduced fitness in fleas [24, 32–35]. The core oligosaccharides of LPS are also occasionally modified with phosphoethanolamine (PEtN) [26, 36]. Similar to Ara4N, PEtN modification reduces negative charge on the LPS [29]. However, this modification predominantly occurs only at very low temperatures (6°C) *in-vitro* and the physiological relevance of this modification is not yet clear. At lower temperatures, lipid A forms containing 3-6 acyl groups are produced, while at temperatures similar to mammalian hosts (37°C), it is predominantly tetraacylated [27]. Although the tetraacylated form may help evade mammalian immune responses [28, 37], it leaves *Y. pestis* more sensitive to the AMP cecropin A [38]. Lipid A acylation-mediated AMP resistance has also been reported in other bacterial species [39, 40]. Transposon mutagenesis studies have demonstrated that AMP resistance in *Y. pestis* is not entirely dependent on LPS modifications, as insertions in genes with no known relationship to LPS-remodeling rendered mutants susceptible to AMPs and reduced survival rates in fleas [32, 41]. Most studies of AMP resistance in *Y. pestis* have used PB as a model peptide [30, 32, 38, 41–43], however, some PB-resistant mutants of *Y. pestis* were not fully resistant to protamine or LL-37, and vice versa, highlighting the fact that resistance to structurally distinct classes of peptides may be different [41]. Therefore, insights into the antimicrobial activities and mechanisms of action of AMPs from natural hosts and vectors are critical to our understanding of AMP resistance and maintenance of wild foci *of Y. pestis.*

Cecropins are one major class of insect antimicrobial peptide with broad spectrum antimicrobial activities [11, 12, 44–47]. In this study, we describe the antimicrobial activity and mechanism of action of a cecropin from *X. cheopis* (cheopin), the first AMP of its class described from Siphonapteran insects, and the role of Ara4N modification in resistance against flea cecropin.

## Materials and Methods

### Synthesis and fluorophore labelling of cheopin

The coding sequence for cecropin was determined from previously published data [19, 48] and verified with draft flea transcriptome data from our laboratory. The amino acid sequence of the mature, active peptide was derived using *in silico* translation and signal peptide detection with *Signal P 5.0* [49]. Insect cecropins are typically amidated at the C-terminus through a post-translational modification by peptidylglycine α-amidating monooxygenase, which removes the glycine and amidates the terminal residue [50, 51]. Therefore, a C-terminal amidated cheopin was chemically synthesized (Biomatik, Wilmington, DE). The purity (>92%) and identity of the peptide was confirmed by reverse phase HPLC and matrix-assisted laser-desorption ionization time of flight (MALDI-TOF) mass spectrometry (calculated 4079.86 /Observed 4078.17). An N-terminal rhodamine labelled cheopin was also chemically synthesized (Biomatik) and a BDP FL-conjugated cheopin was generated by a random labelling bioconjugate method using BDP FL NHS-ester, as recommend by the manufacturer (Lumiprobe, Hunt Valley, MD). Unconjugated fluorophores were removed by ultrafiltration using 3.5 kDa Amicon^®^ centrifugal filters (Sigma-Aldrich, St. Louis, MO). Labelling density was calculated using the correction factor and molar absorption coefficient provided by the manufacturer. A labelling density of 1 dye per peptide was achieved by carefully controlling the reaction stoichiometries (peptide/fluorophore: 1/0.5 molar equivalence).

### Bacterial strains and growth conditions

*Escherichia coli* ATCC^^®^^25922, *Pseudomonas aeruginosa* ATCC^®^27853, *Staphylococcus aureus* ATCC^®^29213, and *Enterococcus faecalis* ATCC^®^29212 were used for antimicrobial susceptibility testing (AST) quality control and to explore the range of antimicrobial activity of cheopin. The details of the *Y. pestis* strains used in this study are given in Table1. Plasmid pMWO-077 was used as a complementation platform [52], and pUC19 [53] was used to express β-lactamase for outer membrane permeability assays. Bacterial cultures except *Y. pestis* were grown in LB or cation-adjusted Mueller-Hinton broth (CAMHB) medium at 37°C, whereas *Y. pestis* were grown in heart infusion medium at 28°C unless otherwise specified (Becton, Dickinson and Company, Franklin Lake, NJ).

The *arn* operon deletion mutant (KIM6+ *ΔarnOP)* was generated in KIM6+ using the λRed recombineering approach [54, 55]. Briefly, a kanamycin resistance gene flanked by FRT sites was amplified from pFKM1 using Q5 high-fidelity polymerase (NEB) and primers with 75 bp oligonucleotide tails homologous to the sequences up and downstream of the *arn* operon (KM9H9-F: ttagttttcgttaacttatctgggcatatagttaatagtccatgaaggtgtcctaagggatttattaatggctcgaattagct-tcaaaag; Y1923_RKM: caagttatatgaatagactaatagggttagtaaataagattagctagcttttataggttttatattaatca-gccaaattggggatcttgaagtacc). The linear PCR product was transformed into KIM6+ electrocompetent cells harboring the pKD46 plasmid [54] and induced with 0.2% arabinose for 4 hours. Recombinants were selected on HI plates with kanamycin and screened for loss of pKD46. Ampicillin-sensitive recombinants were transformed with pFLP2 [56] to remove the Km cassette, leaving a 126 bp in-frame scar site that contained the start codon of the first gene in the operon, *arnB,* as well as the final 12 codons of the last gene, *arnF.* The *arn* operon complementation plasmid was generated by amplifying the entire *arn* operon, including its native promoter region, from KIM6+ with primers arnB_outF (tgctactagtgtatggttgctggctcaagg) and y1923_OutR (tcaaaacgatcacggaatgt) using the AccuStart Taq DNA polymerase HiFi (Quantabio), and cloning it into the *XbaI/PstI* sites of pMWO-077 (a gift from V. Miller, University of North Carolina), replacing the Tet-inducible promoter of pMWO-077. The *arn* operon mutant was then transformed by electroporation with this complementation vector (pMWO77-arnOP) and with an empty vector containing the same promoter deletion (pMWO77). Mutant strains and vectors were sequenced for verification prior to further experimentation.

### Antimicrobial activities

Minimal inhibitory concentrations (MICs), the lowest antimicrobial concentrations at which no visible growth was observed, were determined in CAMH broth using a standardized broth microdilution protocol recommended by the Clinical and Laboratory Standards Institute (CLSI) [58]. MICs were read after 20-24h of incubation, except for *Y. pestis,* which were read after 36h of incubation. Bactericidal concentrations were determined by plating from wells at or above the MIC onto nonselective media, and determining viable colony-forming units (CFU). Assays using *E. coli, P. aeruginosa, S. aureus,* and *E. faecalis* were carried out at 37°C while *Y. pestis* were incubated at 28°C unless otherwise specified.

### CD spectroscopy

Circular dichroism (CD) spectra of the peptides in 10 mM phosphate buffer (pH 7.4) or 10 mM phosphate buffer supplemented with 12 mM SDS were recorded on a Jasco J-815 CD spectrometer. Far UV-spectra (190-250 nm) were recorded with step size of 0.1 nm using a quartz cuvette with path length of 0.1 cm.

### Whole-cell binding assay

The affinity of BDP FL-cheopin for *Y. pestis* was determined by quantifying the fluorescence intensity of peptides bound to bacteria. Briefly, 100μL of cells (1 × 10^7^ CFU/mL in PBS) were incubated with varying concentrations of BDP FL-cheopin for 10 min at room temperature. Cells were then washed twice with 1 mL PBS by centrifugation (5 min, 7000 x g) to remove unbound BDP FL-cheopin. Cell pellets were resuspended in 100 μL of PBS and fluorescence intensities were measured on a BioTek Synergy H1M plate reader (490nm ex, 520 nm em).

### Outer Membrane Permeabilization Assay

The effects of cheopin on the integrity of the bacterial outer membranes were studied by measuring the β-lactamase activity as previously described [59] with slight modifications. Briefly, *Y. pestis* strains carrying the β-lactamase-encoding plasmid pUC19 were grown in the presence of ampicillin to an O.D. of 0.2-0.3, washed twice with 10 mL of PBS-CM (pH 7.4, supplemented with 1 mM CaCl_2_ and 0.5 mM MgCl_2_) by centrifugation (5 min, 5000 x g) and resuspended (1×10^7^ CFU/mL) in PBS-CM containing 180 μM CENTA, a chromogenic β-lactamase substrate (Sigma). Cells (100 μL) were then incubated with 4 μM cheopin or Fast Break lysis solution (Promega, Madison, WI) and absorbance at 405 nm was measured for 1h using a BioTek Synergy H1M plate reader set to 28°C.

### SYTOX Green Uptake Assay

The extent of membrane permeabilization caused by cheopin was evaluated by monitoring intracellular accumulation of the membrane-impermeant dye SYTOX green (Thermo, Waltham, MA). Cells (1×10^7^ CFU/mL) were preincubated with 200 nM SYTOX green in PBS for 10 min at room temperature, treated with 4 μM cheopin or 10% CelLytic B (Sigma), and intracellular accumulation of dye over time was measured by quantifying fluorescence intensity (490 nm ex, 523 nm em) for 30 min on a BioTek Synergy H1M plate reader at 28°C.

### Confocal microscopy

Sub-cellular localization of BDP FL-labelled cheopin was determined using confocal microscopy. Briefly, cells (1×10^8^ CFU/mL) were stained with 1 μM aldehyde-fixable membrane-binding dye FM4-64 FX (Thermo) in PBS for 10 min at room temperature and the unbound dye was removed by washing 2 times with 1 mL PBS by centrifugation (5 min, 7000 x g). Cells were then incubated with 2 μM of BDP FL-cheopin in PBS for 10 min at room temperature and subsequently washed with PBS as in the cell binding assays to remove unbound peptide. Peptide-treated cells were fixed overnight at 4°C in Image-iT fixative solution (Thermo). Fixed cells were washed with 1 mL of PBS (5 min, 10000 x g), and cells were resuspended in 20 μl of ProLong™ glass antifade mountant (Thermo) and spotted on a cover glass. Images were recorded on a Leica SP8 Lightning confocal microscope with a 63X oil immersion objective (NA 1.4). BDPFL and FM4-64 were excited using Argon 488 nm and HeNe 561 nm lasers, with emission signals collected at 510-550 nm and 640-740 nm, respectively. Images were processed for brightness, contrast, and quantification of signal using LAX software (Leica microsystems).

## Results

### X. cheopis cecropin amino acid sequence and characteristics

The mature cheopin peptide sequence was aligned against known cecropin sequences from the Antimicrobial Peptide Database (Fig. 1) [60]. Alignments were performed with ClustalW using the Jukes-Cantor distance model and neighbor-joining algorithm in Geneious prime version 11.1 (Biomatters). The alignment revealed a conserved N-terminal segment, while the C-terminal hydrophobic region showed more variation. Interestingly, cheopin shows substantial similarity to mosquito cecropins, especially in the central and C-terminal regions. Both also have a conserved proline residue in the C-terminal segment, similar to Lepidopteran cecropins, while other cecropins from Diptera do not possess this proline. Although cheopin shows similarity to other known insect cecropins, it is clearly distinct from the sequences identified from Lepidoptera, Diptera, and Coleopteran insects.

**Figure 1.**
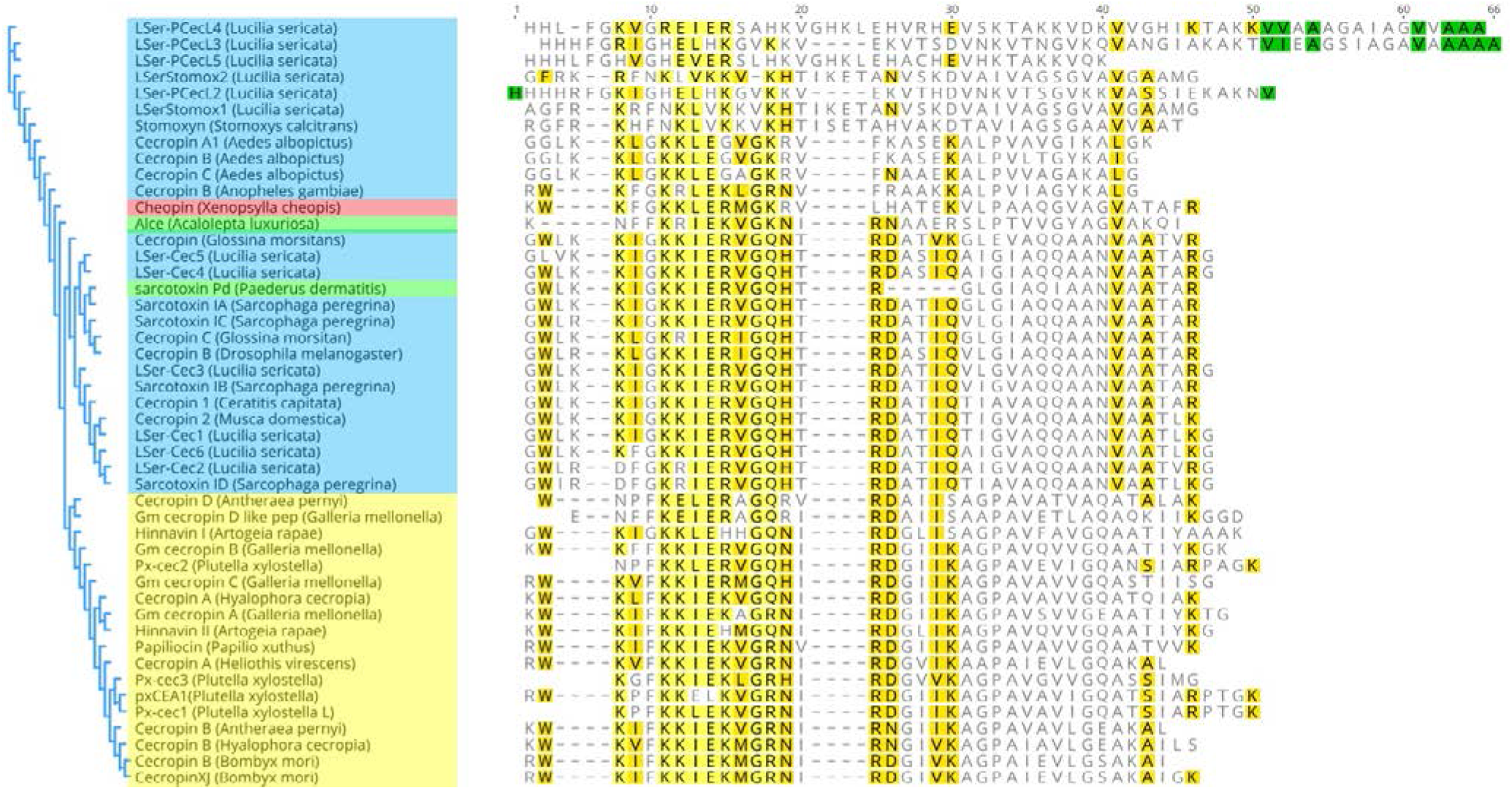
Amino acid sequence of cheopin aligned against known cecropin sequences from different insect orders. Neighbor-joining phylogenetic tree of cecropins from different insect orders. Diptera, Lepidoptera, Coleoptera and Siphonaptera are highlighted in blue, yellow, green and red, respectively. Amino acids colored in green, yellow and gold indicate similarities of 100%, 80-100% and 60-80%, respectively.

### Antimicrobial activities

Antimicrobial susceptibility testing was performed using standardized methods with quality control and *Y. pestis* strains (Table 2). Cheopin showed potent activity against Gram-negative bacteria, but poor activity against Gram-positive organisms. The MICs of cecropin A and cheopin against *E. coli* were comparable, while cheopin was more potent against *P. aeruginosa* compared to cecropin A. Wild-type *Y. pestis* was resistant to all three peptides, including polymyxin B (Table 3). Deletion of the *arn* operon, which is responsible for Ara4N modification of LPS, rendered *Y. pestis* susceptible to both cecropins and polymyxin B. Complementing the *arn* operon mutant restored resistance, confirming the role of Ara4N in AMP resistance in *Y. pestis*.

**Table 1:** *Y. pestis* strains and plasmids used in this study and genotype.

| *Y. pestis* strain or plasmid | Genotype | Reference |
| --- | --- | --- |
| KIM6+ | Parental *Y. pestis* strain; pCD<sup>-</sup>, pMT<sup>+</sup>, pPCP<sup>+</sup> | [57] |
| KIM6+ Δ*arnOP* | In-frame, marker less deletion of *arn* operon (*arnBCADTEF*) in KIM6+ | This study |
| KIM6+ Δ*arnOP* (pMWO77-*arnOP*) | KIM6+ $\Delta$ *arnOP* complemented with pMWO77-*arnOP*, Kan<sup>R</sup>. | This study |
| KIM6 + Δ*arnOP* (pMWO77) | KIM6+ $\Delta$ *arnOP* carrying empty vector pMWO77, Kan<sup>R</sup>. | This study |
| KIM6+ (pUC19) | KIM6+ carrying pUC19 for β-lactamase expression, Amp<sup>R</sup>. | This study |
| KIM6+ Δ*arnOP* (pUC19) | KIM6+ Δ*arnOP* carrying pUC19 for β-lactamase expression, Amp<sup>R</sup>. | This study |
| KIM6+ Δ*arnOP* (pMWO77-*arnOP*, pUC19) | KIM6+ Δ*arnOP* (pMWO77-*arnOP*) carrying pUC19 for β-lactamase expression, Kan<sup>R</sup> Amp<sup>R</sup>. | This study |
| KIM6 + Δ*arnOP* (pMWO77, pUC19) | KIM6+ Δ*arnOP* (pMWO77) carrying pUC19 for β-lactamase expression, Kan<sup>R</sup> Amp<sup>R</sup>. | This study |
| **Plasmids** |  |  |
| pKD46 | λRed recombinase expression plasmid, Amp<sup>R</sup> | [54] |
| pFLP2 | Source of Flp recombinase, Amp<sup>R</sup> | [56] |
| pMWO-077 | Low copy vector for complementation, p15A origin of replication, Kan<sup>R</sup> | [52] |

**Table 2.**
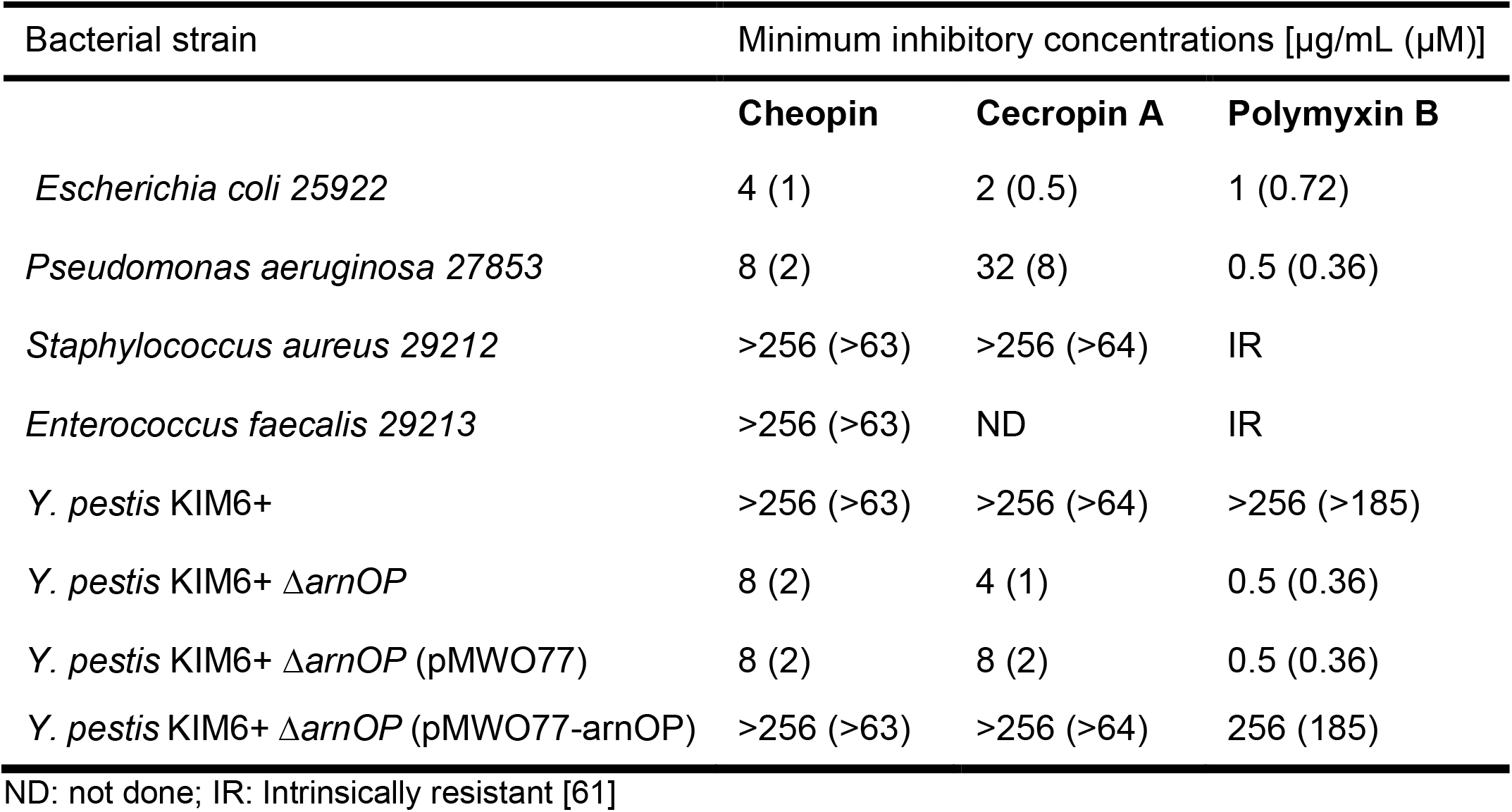
Minimum inhibitory concentrations of cheopin and other AMPs.

### Circular dichroism spectroscopy

CD spectra for cheopin and cecropin A (Fig. 2) show troughs below 200 nm in phosphate buffer, indicating unordered conformation. Characteristic peaks at 196 nm and double troughs at 208 nm and 222 nm indicate that both cecropins adopt α-helical structures in the presence of anionic SDS micelles, similar to other known cecropins [12]. The transition of cheopin to a helical conformation under membrane-like conditions, similar to cecropin A, is consistent with its role as an active cecropin AMP.

**Figure 2:**
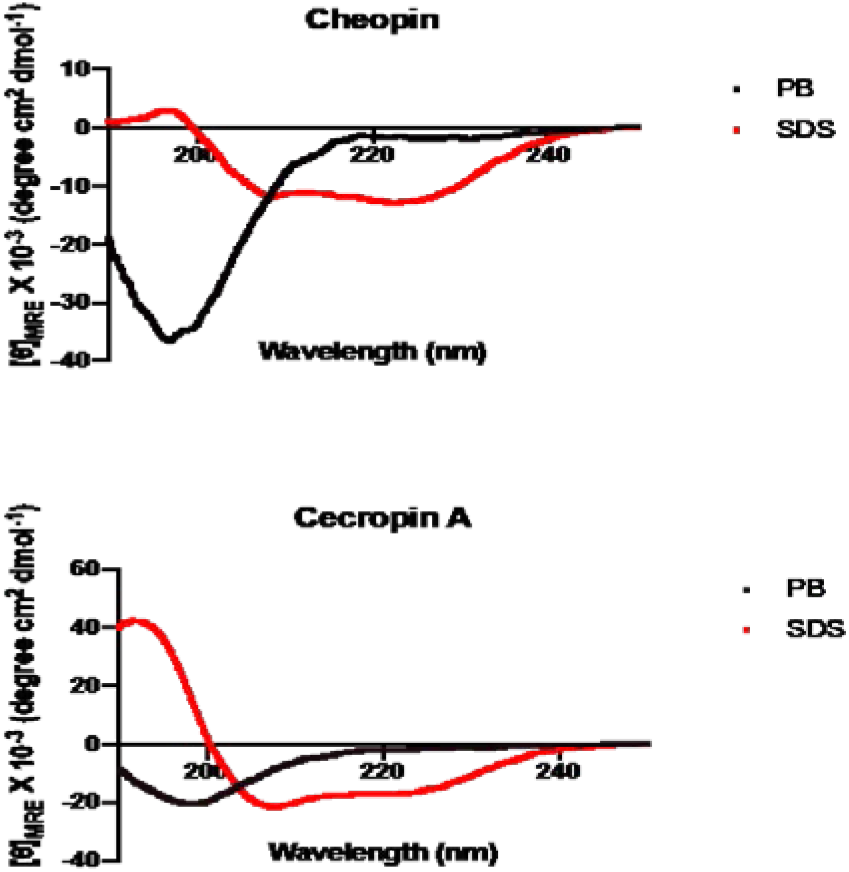
Circular Dichroism spectra of cheopin and cecropin A in 10 mM phosphate buffer (PB) and 12 mM SDS (SDS).

### Whole cell binding assay

To better understand the interaction of cheopin with *Y. pestis*, whole cell binding assays were carried out by measuring the fluorescence remaining after washing cells exposed to fluorophore-labelled cheopin (Fig. 3). Conjugation of rhodamine B to the N-terminus of cheopin led to the loss of antimicrobial activity, therefore, a random labelling method using BDP FL NHS-ester was used to generate fluorophore-conjugated cheopin, which showed the same activity as unlabeled peptide by MIC testing.

**Figure 3:**
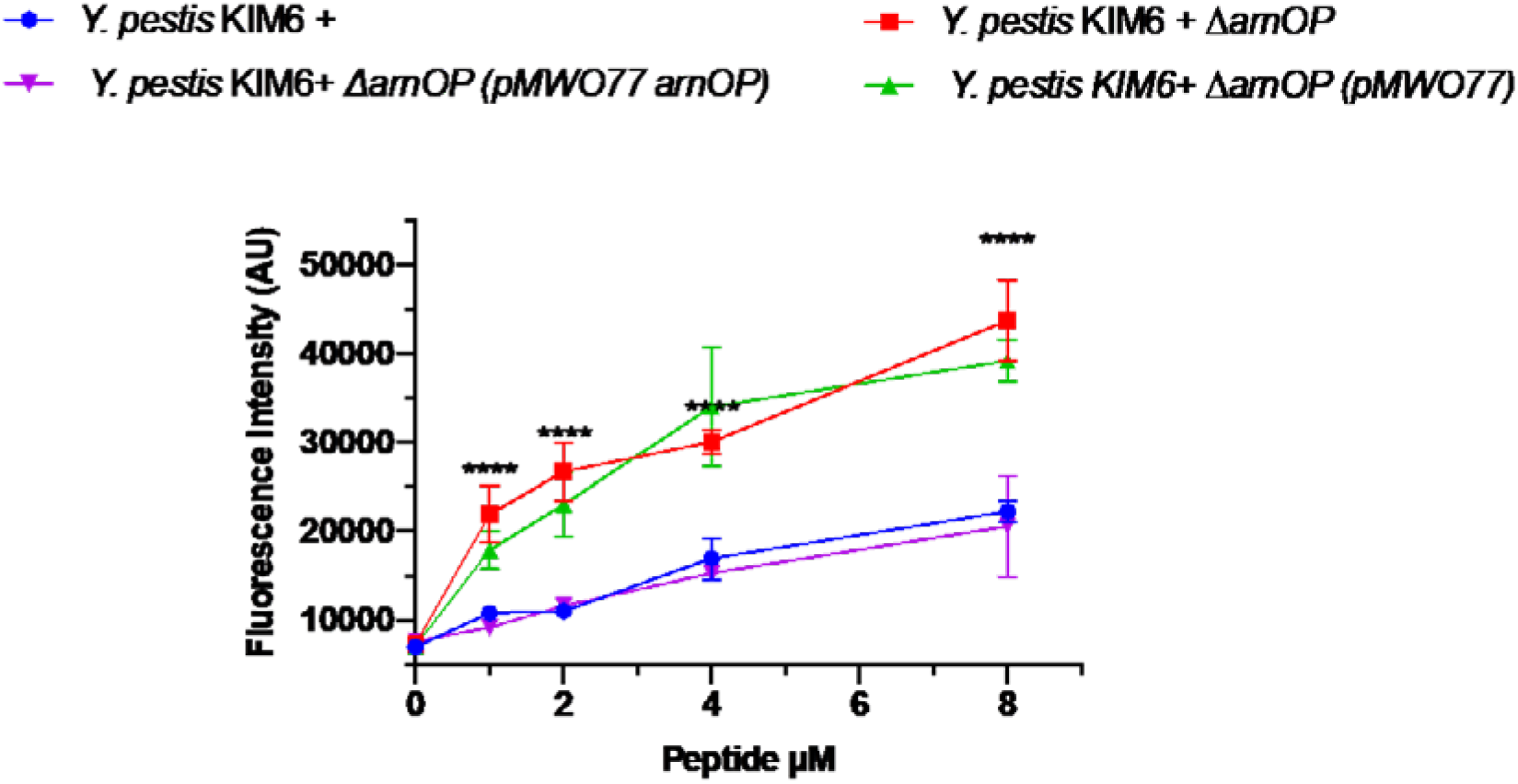
Affinity of BDP FL-cheopin for *Y. pestis* is affected by Ara4N. Cells were treated with BDP FL-labeled cheopin, washed, and fluorescence was acquired to identify cell-associated peptides. Results are the average of three independent experiments and error bars represent standard deviation. Asterisks indicate significant differences (p<0.00001) between wild-type and *arn* operon mutants.

The role of Ara4N modification on binding was determined by comparing wild-type KIM6+ with the Ara4N-deficient *arn* operon deletion mutant. BDP FL-cheopin had significantly higher affinity toward *Y. pestis* KIM6+ *ΔarnOP* than the wild type *Y. pestis* KIM6+, and complementing with a functional *arn* operon reduced the affinity to wild-type levels. Although cheopin exhibited poor antimicrobial activity against wild-type *Y. pestis* (Table 2), it was still able to bind to KIM6+ cells, albeit at significantly lower levels than mutants lacking Ara4N-modified LPS (p<0.00001).

### Effect of cheopin on the integrity of the *Y. pestis* outer membrane

Whole cell binding assays indicated that cheopin binds to wild-type *Y. pestis,* despite its limited activity in MIC testing. Since loss of Ara4N, a positively-charged LPS modification that limits interaction of AMPs with bacterial OMs, increased cheopin binding, we reasoned that bacterial killing is likely to depend on the extent of membrane damage caused by surface-bound peptides. We examined the effect of cheopin on the integrity of the OM by detecting release of the periplasmic enzyme β-lactamase into the extracellular environment with the membrane-impermeant colorimetric substrate CENTA (Fig. 4). Treatment of cells with cheopin did not permeabilize *Y. pestis* KIM6+ OMs, but extensive release of β-lactamase was observed with the *arn* operon mutant and empty vector control. Complementing with the intact *arn* operon restored the integrity of the membrane in the presence of peptide. Clearly, extensive damage of the OM by cheopin is evident only in the absence of Ara4N-modified LPS.

**Figure 4:**
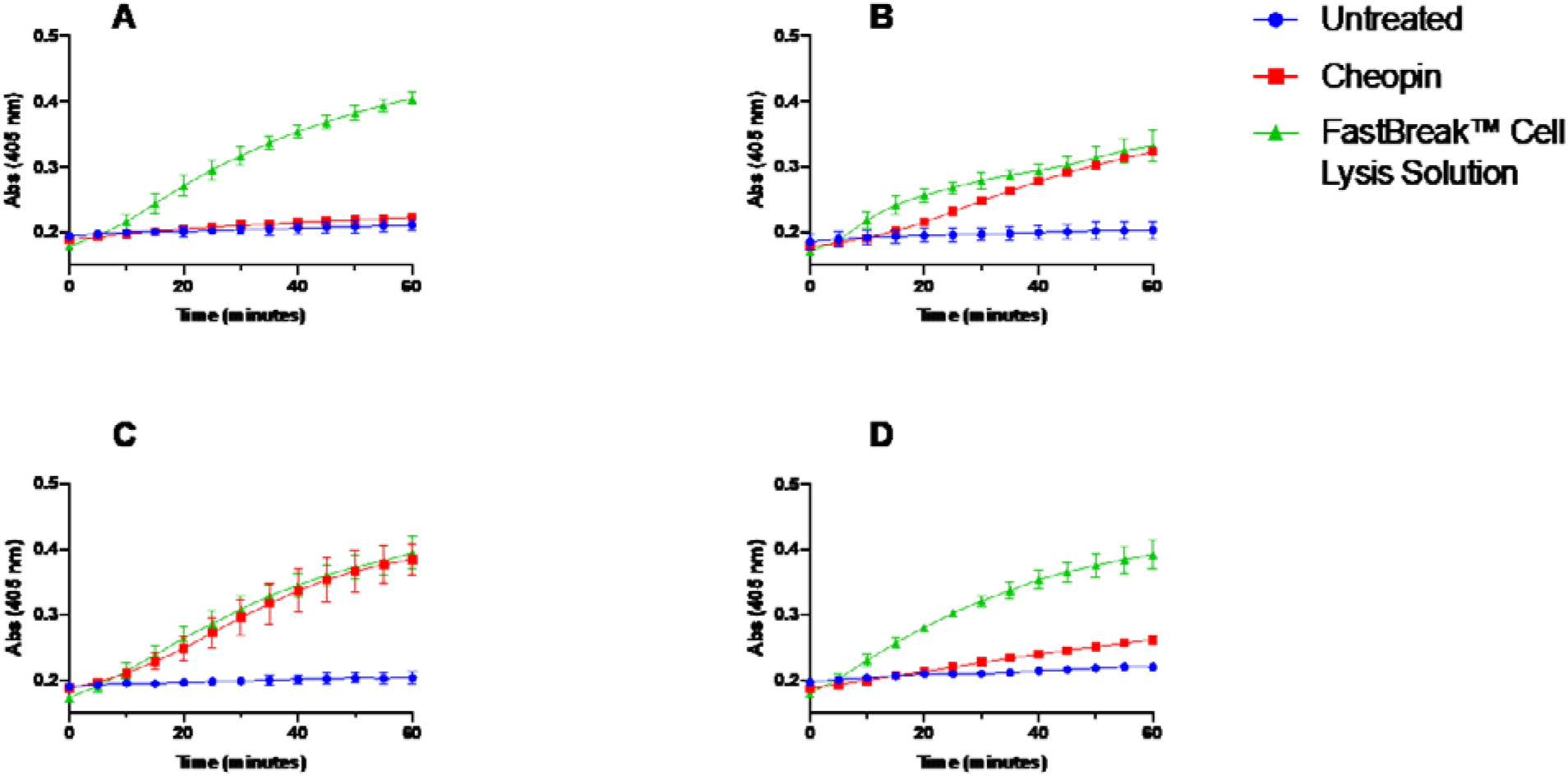
Effect of cheopin on *Y. pestis* outer membranes. The extent of OM damage caused by cheopin was determined by measuring the release of β-lactamase with the colorimetric substrate CENTA. *Y. pestis* strains expressing β-lactamase were treated with 4μM cheopin or FastBreak™ cell lysis solution (positive control) and the hydrolysis of CENTA was monitored for 1h at 28 °C. A: *Y. pestis* KIM6+ (pUC19), B: *Y. pestis KIM6+ΔarnOP* (pUC19), C: *Y. pestis* KIM6+ *ΔarnOP* (pUC19, pMWO77) and D: *Y. pestis* KIM6+ *ΔarnOP* (pUC19, pMWO77-arnOP).

### Effect of cheopin on Y. pestis membranes

To further investigate the mechanism of action, intracellular accumulation of the membrane-impermeant dye SYTOX green was monitored in cheopin-treated cells (Fig. 5). Perturbations of cytoplasmic membranes lead to accumulation of the dye in the cytoplasm, resulting in significantly increased fluorescence upon binding to DNA. Rapid and intense accumulation of SYTOX green was observed in the *arn* operon mutant and empty vector control strain, but only marginal enhancement in fluorescence was observed in the wild-type and complemented strains, again highlighting the important role of Ara4N in limiting the activity of cheopin against *Y. pestis*.

**Figure 5:**
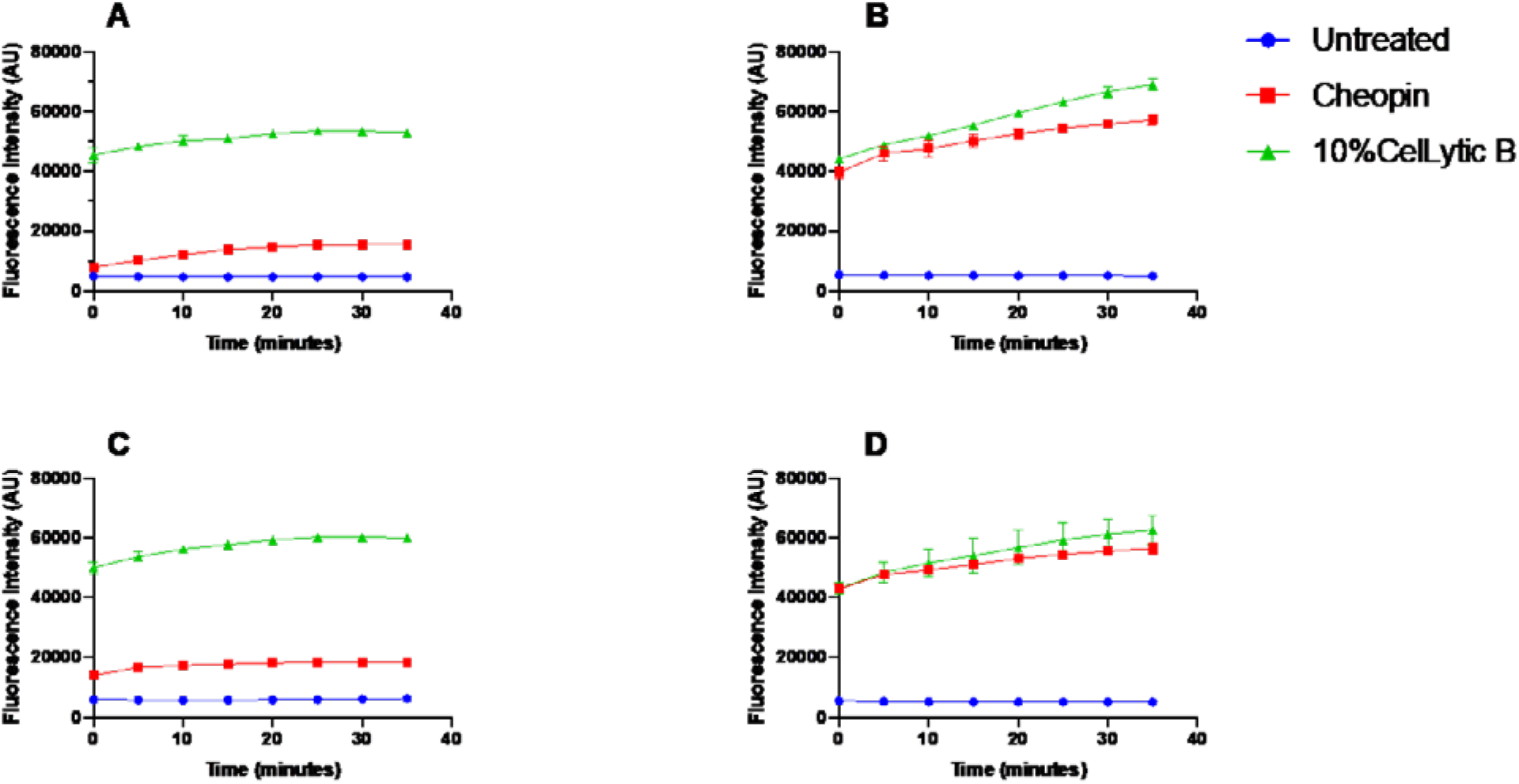
Intracellular accumulation of membrane impermeant dye SYTOX green in cheopin-treated *Y. pestis.* Bacteria were treated with 4 μM cheopin or 10% CelLytic B (positive control) and intracellular accumulation of SYTOX green was measured by monitoring fluorescence. Values are the mean and standard deviations of three independent experiments. A: *Y. pestis* KIM6+, B: *Y. pestis* KIM6+ *ΔarnOP,* C: *Y. pestis* KIM6+ *ΔarnOP* (pMWO77) and D: *Y. pestis* KIM6+ *ΔarnOP* (pMWO77-arnOP).

### Sub-cellular localization of cheopin in Y. pestis

Sub-cellular localization studies using confocal microscopy were carried out to gain further insights into the mechanism of action of cheopin (Fig 6). BDP FL-labeled cheopin showed limited localization to the membranes of KIM6+, as observed with whole cell binding assays, but failed to accumulate in the cytoplasm. The absence of Ara4N in the *arn* operon mutant and empty vector control led to more intense binding of peptide to the membranes and allowed penetration of cheopin into the cytoplasm. Some mutant cells showed limited intracellular signal from the peptide, likely due to asynchronous kinetics of binding and penetration of cells. Complementing the deleted *arn* operon reduced the localization of peptide to the cells, illustrating the protective effect of Ara4N on cheopin binding and intracellular penetration.

**Figure 6:**
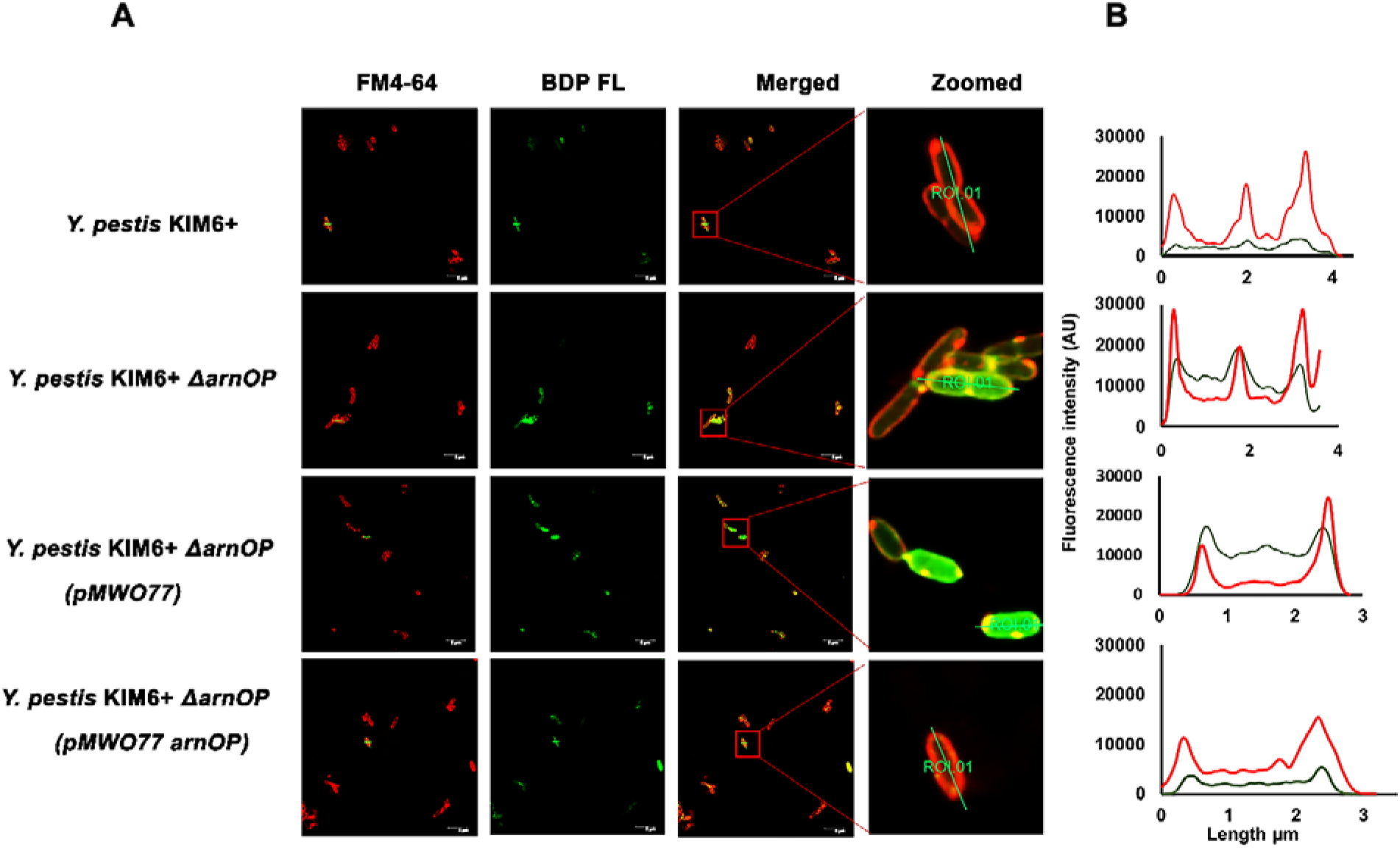
Sub-cellular localization of cheopin in *Y. pestis.* Localization of BDP FL-cheopin (green) on *Y. pestis* pre-stained with the cytoplasmic membrane dye FM4-64 FX (red). The images in the far-right column are zoomed in at the locations shown in the merged images to more clearly illustrate co-localization (Fig. 6A). The graphs represent fluorescence intensities across the selected regions of interest (ROI, Fig. 6B). Scale bars 5 μm.

## Discussion

Cecropins are one of the major groups of linear cationic peptides produced by insects. Cecropins have been identified in many insects from the Orders Diptera, Lepidoptera and Coleoptera [11, 12, 44–47, 50, 62]. In this study, we describe the antimicrobial activities and mechanism of action of a novel cecropin, cheopin, from *X. cheopis* of the order Siphonaptera. The amino acid sequence of cheopin shows remarkable similarities to other known cecropins, yet with distinct features. While the N-terminal amphipathic segment of cheopin is similar to other insect cecropins, the C-terminal region shows considerable differences. Intriguingly, cheopin shows substantial similarity to mosquito cecropins and has a highly conserved proline residue following the N-terminal amphipathic segment, similar to Lepidopteran cecropins. Both mosquitos and fleas share many common hosts including humans: consequently, exposure to similar microorganisms is likely. It is plausible that common host-factors such as this may have played a role in the selection of cecropin sequences.

Insect cecropins are known to possess broad spectrum antimicrobial activity against Gramnegative and Gram-positive bacteria, fungi and parasites. However, potencies toward Gramnegative bacteria are several fold higher than against Gram-positive organisms [12, 44, 46, 47, 62, 63]. Cheopin, similar to the well-studied cecropin A, follows this pattern, with potent activity against Gram-negative bacteria such as *E. coli* and *P. aeruginosa,* but little activity against the Gram-positive organisms tested. Structure-activity relationship studies on cecropins have identified that the N-terminal amphipathic segment is important for antimicrobial activity against many organisms [62, 64], and subtle variations in insect cecropin sequences are known to play a role in antibacterial activity. Replacement of a highly conserved tryptophan residue with glycine in the N-terminal segment of a mosquito cecropin was essential for the activity against Gram-positive bacteria [47]. Similarly, deletion of C-terminal residues reduced the activity of cecropin A against *Micrococcus luteus* and *Bacillus megaterium,* but not against *E. coli* or other bacteria [62, 65]. Although the cheopin sequence is considerably different from cecropin A, many of the differences are in the C-terminal hydrophobic segment. Therefore, as seen in other cecropins, these C-terminal differences may be responsible for the 4 fold lower MIC against *P. aeruginosa* for cheopin vs. cecropin A.

Though cheopin was active against *E. coli* and *P. aeruginosa,* it showed little activity against wild-type *Y. pestis* KIM6+ under conditions promoting Ara4N modification of LPS. However, when the *arn* operon was deleted, *Y. pestis* was rendered highly susceptible to this novel AMP. Complementation with an intact *arn* operon restored resistance, demonstrating the critical role of Ara4N-modified LPS in protecting *Y. pestis* against cheopin. This is consistent with previous observations that mutants lacking Ara4N were significantly attenuated for survival in fleas [32].

The bactericidal mechanism of cecropin and many other cationic AMPs involves sequential permeabilization of outer and inner membranes [39, 66–70]. Confocal microscopy revealed that cheopin localized to *Y. pestis* membranes and cytoplasm, the latter indicating an ability to disrupt both membranes. Interestingly, cheopin bound to *Y. pestis* KIM6+ despite its resistance to killing, albeit to a lesser extent than the mutant lacking Ara4N-modified LPS. While confocal microscopy and whole cell binding assays indicated that cheopin binds to *Y. pestis,* extensive permeabilization of outer and inner membranes was observed only in the absence of Ara4N-modified LPS. This is consistent with the idea that Ara4N-mediated charge shielding of the outer membranes allows *Y. pestis* KIM6+ to successfully evade the assault by cheopin. Time-lapse microscopy studies with cecropin A revealed a two-step process for permeabilization of bacterial membranes: an initial, slow interaction of peptide with cell surfaces, followed by a more rapid nucleation event after interaction with charged lipid A head groups. This nucleation event leads to OM breach and subsequent, inevitable disruption of the cytoplasmic membrane [68, 69]. The killing mediated by cheopin appears to behave in the same manner, with permeabilization of the outer membrane, destabilization of the cytoplasmic membrane and localization into the cytoplasm.

In this study, we report membrane permeabilization and bactericidal activity of the first characterized Siphonapteran cecropin. Cheopin kills bacteria like other cecropins, by permeabilizing membranes, and possibly interfering with cytoplasmic processes. Further, our results provide direct evidence for the role of Ara4N modification of LPS by *Y. pestis* in resisting an antimicrobial peptide from its flea vector.

## Acknowledgments

This work was supported by a grant (R01AI130255) from the National Institutes of Allergy and Infectious Diseases. We thank the University of Utah Health science Imaging, DNA and Peptide, and Proteomics core facilities for their help with confocal microscopy, HPLC and Mass spectrometry. We also acknowledge the University of Utah Optical Spectroscopy core for their help with the CD-spectroscope.

## Notes

### Competing Interest Statement

The authors have declared no competing interest.

## References

1. Nikiforov, V.V., et al., Plague: Clinics, Diagnosis and Treatment. Adv Exp Med Biol, 2016. 918: p. 293–312.

2. Bi, Y., Immunology of Yersinia pestis Infection. Adv Exp Med Biol, 2016. 918: p. 273–292.

3. Shi, L., et al., Reemergence of human plague in Yunnan, China in 2016. PLoS One, 2018. 13(6): p. e0198067.

4. Abedi, A.A., et al., Ecologic Features of Plague Outbreak Areas, Democratic Republic of the Congo, 2004-2014. Emerg Infect Dis, 2018. 24(2): p. 210–220.

5. Andrianaivoarimanana, V., et al., Trends of Human Plague, Madagascar, 1998-2016. Emerg Infect Dis, 2019. 25(2): p. 220–228.

6. Demeure, C.E., et al., Yersinia pestis and plague: an updated view on evolution, virulence determinants, immune subversion, vaccination, and diagnostics. Genes Immun, 2019. 20(5): p. 357–370.

7. Galimand, M., E. Carniel, and P. Courvalin, Resistance of Yersinia pestis to antimicrobial agents. Antimicrob Agents Chemother, 2006. 50(10): p. 3233–6.

8. Dubyanskiy, V.M. and A.B. Yeszhanov, Ecology of Yersinia pestis and the Epidemiology of Plague, in Yersinia pestis: Retrospective and Perspective, R. Yang and A. Anisimov, Editors. 2016, Springer Netherlands: Dordrecht. p. 101–170.

9. Mullen, G.R., L.A. Durden, and G. Mullen, Medical and Veterinary Entomology. 2002, Burlington, UNITED STATES: Elsevier Science & Technology.

10. Brown, L.D., Immunity of fleas (Order Siphonaptera). Dev Comp Immunol, 2019. 98: p. 76–79.

11. Wu, Q., J. Patocka, and K. Kuca, Insect Antimicrobial Peptides, a Mini Review. Toxins (Basel), 2018. 10(11).

12. Yi, H.Y., et al., Insect antimicrobial peptides and their applications. Appl Microbiol Biotechnol, 2014. 98(13): p. 5807–22.

13. Feldhaar, H. and R. Gross, Immune reactions of insects on bacterial pathogens and mutualists. Microbes Infect, 2008. 10(9): p. 1082–8.

14. Kartman, L. and F.M. Prince, Studies on Pasteurella pestis in fleas. V. The experimental plague-vector efficiency of wild rodent fleas compared with Xenopsylla cheopis, together with observations on the influence of temperature. Am J Trop Med Hyg, 1956. 5(6): p. 1058–70.

15. Kartman, L., F.M. Prince, and S.F. Quan, Studies on Pasteurella pestis in fleas, comparative plague-vector efficiency of Xenopsylla vexabilis hawaiiensis and Xenopsylla cheopis. Bull World Health Organ, 1956. 14(4): p. 681–704.

16. Gage, K.L. and M.Y. Kosoy, Natural history of plague: perspectives from more than a century of research. Annu Rev Entomol, 2005. 50: p. 505–28.

17. Andersen, J.F., et al., An insight into the sialome of the oriental rat flea, Xenopsylla cheopis (Rots). BMC Genomics, 2007. 8: p. 102.

18. Bland, D.M., et al., Transcriptomic profiling of the digestive tract of the rat flea, Xenopsylla cheopis, following blood feeding and infection with Yersinia pestis. PLOS Neglected Tropical Diseases, 2020. 14(9): p. e0008688.

19. Consortium, G.R.D., et al., Genomic Resources Notes Accepted 1 August 2015 – 31 September 2015. 2016. 16(1): p. 377–377.

20. Hinnebusch, B.J., C.O. Jarrett, and D.M. Bland, “Fleaing” the Plague: Adaptations of Yersinia pestis to Its Insect Vector That Lead to Transmission. Annu Rev Microbiol, 2017. 71: p. 215–232.

21. Vadyvaloo, V., et al., Role of the PhoP-PhoQ gene regulatory system in adaptation of Yersinia pestis to environmental stress in the flea digestive tract. Microbiology, 2015. 161(6): p. 1198–1210.

22. Zhou, D., et al., Transcriptome analysis of the Mg2+-responsive PhoP regulator in Yersinia pestis. FEMS Microbiol Lett, 2005. 250(1): p. 85–95.

23. Han, Y., et al., Comparative transcriptome analysis of Yersinia pestis in response to hyperosmotic and high-salinity stress. Res Microbiol, 2005. 156(3): p. 403–15.

24. Vadyvaloo, V., et al., Transit through the flea vector induces a pretransmission innate immunity resistance phenotype in Yersinia pestis. PLoS Pathog, 2010. 6(2): p. e1000783.

25. Anisimov, A.P., et al., Intraspecies and temperature-dependent variations in susceptibility of Yersinia pestis to the bactericidal action of serum and to polymyxin B. Infect Immun, 2005. 73(11): p. 7324–31.

26. Knirel, Y.A. and A.P. Anisimov, Lipopolysaccharide of Yersinia pestis, the Cause of Plague: Structure, Genetics, Biological Properties. Acta Naturae, 2012. 4(3): p. 46–58.

27. Knirel, Y.A., et al., Temperature-dependent variations and intraspecies diversity of the structure of the lipopolysaccharide of Yersinia pestis. Biochemistry, 2005. 44(5): p. 1731–43.

28. Rebeil, R., et al., Variation in lipid A structure in the pathogenic yersiniae. Mol Microbiol, 2004. 52(5): p. 1363–73.

29. Band, V.I. and D.S. Weiss, Mechanisms of Antimicrobial Peptide Resistance in Gram-Negative Bacteria. Antibiotics (Basel), 2015. 4(1): p. 18–41.

30. Rebeil, R., et al., Induction of the Yersinia pestis PhoP-PhoQ regulatory system in the flea and its role in producing a transmissible infection. J Bacteriol, 2013. 195(9): p. 1920–30.

31. Hitchen, P.G., et al., Structural characterization of lipo-oligosaccharide (LOS) from Yersinia pestis: regulation of LOS structure by the PhoPQ system. Mol Microbiol, 2002. 44(6): p. 1637–50.

32. Aoyagi, K.L., et al., LPS modification promotes maintenance of Yersinia pestis in fleas. Microbiology, 2015. 161(Pt 3): p. 628–38.

33. Earl, S.C., et al., Resistance to Innate Immunity Contributes to Colonization of the Insect Gut by Yersinia pestis. PLoS One, 2015. 10(7): p. e0133318.

34. Erickson, D.L., et al., PhoP and OxyR transcriptional regulators contribute to Yersinia pestis virulence and survival within Galleria mellonella. Microb Pathog, 2011. 51(6): p. 389–95.

35. Fukuto, H.S., et al., A Single Amino Acid Change in the Response Regulator PhoP, Acquired during Yersinia pestis Evolution, Affects PhoP Target Gene Transcription and Polymyxin B Susceptibility. J Bacteriol, 2018. 200(9).

36. Knirel, Y.A., et al., Cold temperature-induced modifications to the composition and structure of the lipopolysaccharide of Yersinia pestis. Carbohydr Res, 2005. 340(9): p. 1625–30.

37. Montminy, S.W., et al., Virulence factors of Yersinia pestis are overcome by a strong lipopolysaccharide response. Nat Immunol, 2006. 7(10): p. 1066–73.

38. Rebeil, R., et al., Characterization of late acyltransferase genes of Yersinia pestis and their role in temperature-dependent lipid A variation. J Bacteriol, 2006. 188(4): p. 1381–8.

39. Guo, L., et al., Lipid A acylation and bacterial resistance against vertebrate antimicrobial peptides. Cell, 1998. 95(2): p. 189–98.

40. Joo, H.S., C.I. Fu, and M. Otto, Bacterial strategies of resistance to antimicrobial peptides. Philos Trans R Soc Lond B Biol Sci, 2016. 371(1695).

41. Guo, J., et al., Tn5AraOut mutagenesis for the identification of Yersinia pestis genes involved in resistance towards cationic antimicrobial peptides. Microb Pathog, 2011. 51(3): p. 121–32.

42. Bengoechea, J.A., et al., Yersinia pseudotuberculosis and Yersinia pestis are more resistant to bactericidal cationic peptides than Yersinia enterocolitica. Microbiology, 1998. 144 (Pt 6): p. 1509-15.

43. Zhou, D., et al., Genome-wide transcriptional response of Yersinia pestis to stressful conditions simulating phagolysosomal environments. Microbes Infect, 2006. 8(12-13): p. 2669–78.

44. Steiner, H., et al., Sequence and specificity of two antibacterial proteins involved in insect immunity. Nature, 1981. 292(5820): p. 246–8.

45. Hultmark, D., et al., Insect immunity: isolation and structure of cecropin D and four minor antibacterial components from Cecropia pupae. Eur J Biochem, 1982. 127(1): p. 207–17.

46. Lowenberger, C., et al., Antimicrobial activity spectrum, cDNA cloning, and mRNA expression of a newly isolated member of the cecropin family from the mosquito vector Aedes aegypti. J Biol Chem, 1999. 274(29): p. 20092–7.

47. Vizioli, J., et al., Cloning and analysis of a cecropin gene from the malaria vector mosquito, Anopheles gambiae. Insect Mol Biol, 2000. 9(1): p. 75–84.

48. Driver, J.D., The antibacterial immune response to Escherichia coli in the flea Xenopsylla cheopis. 2002, https://scholarworks.umt.edu/etd/9385

49. Almagro Armenteros, J.J., et al., SignalP 5.0 improves signal peptide predictions using deep neural networks. Nature Biotechnology, 2019. 37(4): p. 420–423.

50. Lidholm, D.-A., et al., Insect immunity: cDNA clones coding for the precursor forms of cecropins A and D, antibacterial proteins from Hyalophora cecropia. 1987. 226(1): p. 8–12.

51. Vishwanatha, K.S., R.E. Mains, and B.A. Eipper, *Chapter 244 - Peptidylglycine Amidating Monoxygenase (PAM)*, in Handbook of Biologically Active Peptides (Second Edition), A.J. Kastin, Editor. 2013, Academic Press: Boston. p. 1780–1788.

52. Obrist, M.W. and V.L. Miller, Low copy expression vectors for use in Yersinia sp. and related organisms. Plasmid, 2012. 68(1): p. 33–42.

53. Yanisch-Perron, C., J. Vieira, and J. Messing, Improved M13 phage cloning vectors and host strains: nucleotide sequences of the M13mpl8 and pUC19 vectors. Gene, 1985. 33(1): p. 103–119.

54. Datsenko, K.A. and B.L. Wanner, One-step inactivation of chromosomal genes in Escherichia coli K-12 using PCR products. Proc Natl Acad Sci U S A, 2000. 97(12): p. 6640–5.

55. Derbise, A., et al., A rapid and simple method for inactivating chromosomal genes in Yersinia. FEMS Immunol Med Microbiol, 2003. 38(2): p. 113–6.

56. Hoang, T.T., et al., A broad-host-range Flp-FRT recombination system for site-specific excision of chromosomally-located DNA sequences: application for isolation of unmarked Pseudomonas aeruginosa mutants. Gene, 1998. 212(1): p. 77–86.

57. Perry, R.D., M.L. Pendrak, and P. Schuetze, Identification and cloning of a hemin storage locus involved in the pigmentation phenotype of Yersinia pestis. J Bacteriol, 1990. 172(10): p. 5929–37.

58. Wayne, P., ed. CLSI. Methods for Dilution Antimicrobial Susceptibilty Tests for Bacteria That Grow Aerobically. 11th ed. CLSI standard M07. 2018.

59. Luther, A., et al., Chimeric peptidomimetic antibiotics against Gram-negative bacteria. Nature, 2019. 576(7787): p. 452–458.

60. Wang, G., X. Li, and Z. Wang, APD3: the antimicrobial peptide database as a tool for research and education. Nucleic Acids Res, 2016. 44(D1): p. D1087–93.

61. Wayne, P., ed. CLSI. Methods for Dilution Antimicrobial Susceptibilty Tests for Bacteria That Grow Aerobically. 11th ed. CLSI standard M100. 2018.

62. Andreu, D., et al., N-terminal analogues of cecropin A: synthesis, antibacterial activity, and conformational properties. Biochemistry, 1985. 24(7): p. 1683–8.

63. Kim, W., et al., Ectopic expression of a cecropin transgene in the human malaria vector mosquito Anopheles gambiae (Diptera: Culicidae): effects on susceptibility to Plasmodium. J Med Entomol, 2004. 41(3): p. 447–55.

64. Lee, E., et al., Functional Roles of Aromatic Residues and Helices of Papiliocin in its Antimicrobial and Anti-inflammatory Activities. Scientific Reports, 2015. 5(1): p. 12048.

65. Merrifield, R.B., L.D. Vizioli, and H.G. Boman, Synthesis of the antibacterial peptide cecropin A (1-33). Biochemistry, 1982. 21(20): p. 5020–31.

66. Brogden, K.A., Antimicrobial peptides: pore formers or metabolic inhibitors in bacteria? Nat Rev Microbiol, 2005. 3(3): p. 238–50.

67. Lee, T.H., K.N. Hall, and M.I. Aguilar, Antimicrobial Peptide Structure and Mechanism of Action: A Focus on the Role of Membrane Structure. Curr Top Med Chem, 2016. 16(1): p. 25–39.

68. Rangarajan, N., S. Bakshi, and J.C. Weisshaar, Localized permeabilization of E. coli membranes by the antimicrobial peptide Cecropin A. Biochemistry, 2013. 52(38): p. 6584–94.

69. Agrawal, A. and J.C. Weisshaar, Effects of alterations of the E. coli lipopolysaccharide layer on membrane permeabilization events induced by Cecropin A. Biochim Biophys Acta Biomembr, 2018. 1860(7): p. 1470–1479.

70. Silvestro, L., J.N. Weiser, and P.H. Axelsen, Antibacterial and antimembrane activities of cecropin A in Escherichia coli. Antimicrob Agents Chemother, 2000. 44(3): p. 602–7.

